# Responses of Neurons in the Rostral Ventrolateral Medulla (RVLM) of Conscious Felines to Anticipated and Passive Movements

**DOI:** 10.1101/693408

**Authors:** Derek M. Miller, Asmita Joshi, Emmanuel T. Kambouroglos, Isaiah C. Engstrom, John P. Bielanin, Samuel R. Wittman, Andrew A. McCall, Susan M. Barman, Bill J. Yates

## Abstract

Considerable evidence demonstrates that the vestibular system contributes to regulating sympathetic nerve activity and blood pressure. Initial studies in decerebrate animals showed that presumed pre-sympathetic neurons in the rostral ventrolateral medulla (RVLM) respond to small-amplitude (<10°) rotations of the body, as in other brain areas that process vestibular signals, despite the fact that such movements do not appreciably affect blood distribution in the body. However, a subsequent experiment in conscious animals showed that few RVLM neurons respond to small-amplitude movements. This study tested the hypothesis that vestibular inputs to RVLM neurons are modulated in conscious animals, such that vestibulosympathetic responses are only elicited when changes in body position are large enough to require changes in sympathetic nerve activity. The activity of approximately a third of RVLM neurons whose firing rate was related to the cardiac cycle, and thus likely received baroreceptor inputs, responded to vestibular inputs elicited by 40° head-up tilts in conscious cats, but not during 10° sinusoidal rotations in the pitch plane that affected the activity of neurons in brain regions providing inputs to the RVLM. These data suggest the existence of brain circuitry that suppresses vestibular influences on the activity of RVLM neurons and the sympathetic nervous system unless these inputs are physiologically warranted. We also determined that RVLM activity is not altered prior to tilts when a light cue is provided signaling the movement. The simplest interpretation of this findings is that feedforward cardiovascular responses are associated with active movement such as occurs during exercise, but not passive movements that require cardiovascular adjustments.

## INTRODUCTION

When either humans or quadrupeds transition from supine or prone to a head-up position (as during standing in humans), gravity causes an increase in arterial and venous pressures below the heart and a decrease in pressures above the heart (62). As a consequence, there is an increase in lower body intravascular fluid volume and a decrease in venous return to the heart, which can result in reduced cardiac output and blood flow to the brain (7, 20, 21, 60). Orthostatic hypotension occurs following these physiologic alterations unless there are rapid compensatory changes in sympathetic nerve activity (SNA) (7).

It has long been appreciated that unloading of baroreceptors and perhaps cardiac mechanoreceptors during postural changes results in increased SNA (6, 52). In addition, there is now substantial evidence from both human and animal subjects that the vestibular system contributes to maintaining stable blood pressure and cerebral blood flow during movement and postural alterations (62). Conversely, loss of vestibular inputs diminishes compensatory changes in SNA during postural changes, increasing the likelihood that orthostatic hypotension will occur (26, 61).

Several studies have explored the neural circuitry that produces vestibulosympathetic responses (VSR). Anatomical studies in rodents demonstrate direct projections from the caudal aspect of the vestibular nuclei to the rostral ventrolateral medulla (RVLM) (23, 24), which contains reticulospinal neurons that regulate SNA (33, 51, 56). In addition, lesions of the RVLM abolish changes in SNA elicited by electrical stimulation of vestibular afferent fibers (55). Studies in decerebrate felines show that RVLM neurons respond robustly to electrical stimulation of vestibular afferents (64) or small-amplitude (7.5°-10°) rotations of the head (12, 63), supporting the notion that RVLM neurons transmit vestibular signals to sympathetic preganglionic neurons in the thoracic spinal cord.

In more recent studies, the activity of RVLM neurons was recorded in conscious felines as their body was passively rotated (12). Unlike in decerebrate animals, virtually no RVLM neurons in conscious cats responded to small-amplitude rotations that strongly modulate the activity of neurons in a number of brain regions, including the cerebellar fastigial nucleus (43), vestibular nuclei (37), and lateral tegmental field (39). These findings appeared to contradict the notion that RVLM neurons play a key role in mediating VSR. However, it was also noted that head-up body rotations less than 40° in amplitude do not result in fluid shifts that can appreciably affect cardiac output (65). These observations raised the hypothesis that vestibular signals are integrated in the VSR circuit such that changes in sympathetic outflow are only triggered by large, sudden head-up postural alterations with the potential of producing orthostatic hypotension (38). Furthermore, this integrative mechanism is affected by decerebration, such that RVLM neurons respond to smaller changes in head position than in conscious animals (12).

As a first step to explore this possibility, in the current study the activity of RVLM neurons was recorded during 40° trapezoidal (ramp-and-hold) head-up pitch rotations in conscious cats. For many units, we also examined responses to 10° peak-to-peak sinusoidal rotations in the pitch plane similar to those used in prior studies. We tested the hypothesis that a large fraction of RVLM neurons respond to large amplitude pitch rotations, but not to smaller amplitude rotations that robustly modulate activity of neurons in other brain areas that process vestibular inputs.

An extensive literature shows that cardiovascular responses can precede the onset of exercise (13, 16, 59). However, it is unclear whether feedforward cardiovascular responses are initiated prior to other movements that require coupled changes in SNA, such as head-up postural alterations. Cardiovascular responses to a variety of stimuli can be conditioned in animals and humans, such that they are subsequently elicited by cues that are associated with the triggering event (34, 41). In a prior study, a light cue preceded 60° head-up rotations of conscious felines that were instrumented to record cerebral blood flow, to ascertain whether brain perfusion or heart rate were modified by a conditioning stimulus paired to the postural alteration (46). This study found no evidence that either heart rate or cerebral blood flow were altered after the cue and prior to the onset of body movement. In addition, variations in heart rate and cerebral blood flow were not noted in a subset of trials in which the anticipated head-up tilt was not provided after the cue. A criticism of this study was that it failed to consider a variety of cardiovascular responses elicited during head-up tilts such as vasoconstriction in the lower body. Since RVLM neurons play a key role in regulating vascular smooth muscle contraction mediated by the sympathetic nervous system (33, 51, 56), an examination of RVLM activity prior to and during cued head-up rotations is potentially a more definitive test of whether feedforward cardiovascular responses precede expected passive postural alterations that affect blood distribution in the body. Accordingly, these experiments tested a second hypothesis that RVLM neuronal activity is modified prior to the onset of 40° head-up tilts signaled by a prominent visual cue.

Like most areas of the reticular formation, the RVLM is heterogeneous (18, 33, 51, 56). In addition to using postmortem histological reconstructions to distinguish which sampled units were located anatomically in the RVLM, our analyses focused on the subset of units with cardiac-related activity (CRA), or rhythmic changes in activity correlated with cardiac contractions. The presence of CRA is a hallmark indicator that a brainstem neuron participates in the regulation of SNA (5).

## METHODS

All experimental procedures conformed to the American Physiological Society’s “Guiding Principles for the Care and Use of Animals,” as well as the National Research Council *Guide for the Care and Use of Laboratory Animals*, and were approved by the University of Pittsburgh’s Institutional Animal Care and Use Committee. Experiments were conducted over a period of ~7 months on two purpose-bred adult female cats (Liberty Research, Waverly, NY). Use of two animals in studies entailing the sampling of neuronal activity from a large number of neurons over time is typical in neurophysiological studies (22, 27, 45, 50, 54, 58).

Experimental procedures in this study were similar to those in our prior experiments (4, 12, 37, 39, 40, 42, 43) and entailed acclimating animals for body restraint, implanting instrumentation to record single unit activity from individual brainstem neurons as well as the electrocardiogram (ECG) and carotid artery blood flow, another period of acclimation for restraint, and a period of 4-5 months during which recordings of activity of brainstem neurons were obtained. At the end of experiments, animals were euthanized and the locations of recording sites were histologically reconstructed.

### Surgical Procedures

Animals were spayed by a veterinarian prior to the onset of studies to eliminate cyclic changes in hormonal levels that could affect physiologic responses. Subsequently, over a period of ~2 months, animals were gradually acclimated for body restraint. Initially, animals were restrained for only a few minutes per session; over subsequent training sessions the restraint period was gradually increased to a 2-hour period.

After animals were acclimated for restraint, a recovery surgery was performed using aseptic techniques in a dedicated operating room that entailed a craniectomy, attaching a recording chamber and head fixation plate to the skull, and implanting subcutaneous ECG electrodes and perivascular probes (PS series, Transonic Systems, Ithaca, NY) around the common carotid arteries. These surgical procedures have been described in detail (4, 12). Anesthesia was induced with an intramuscular injection of ketamine (20 mg/kg) and acepromazine (0.2 mg/kg), an endotracheal tube was inserted, and anesthesia was subsequently maintained using 1-2% isoflurane vaporized in oxygen. After surgeries were completed, analgesia was provided for 72 hrs through transdermal administration of 25 μg/hr fentanyl. In addition, antibiotics (amoxicillin, two 50-mg oral doses per day) were provided for 10 days.

### Recording Procedures

Animals recovered from surgery for at least 2-weeks prior to the resumption of experimental procedures. The restraint device securing the animal’s body was mounted on a computer-controlled hydraulic tilt table (Neurokinetics, Pittsburgh, PA). Over a period of ~ 1 month, animals were acclimated for head restraint by inserting a screw into the head-mounted fixation plate, as well as to whole-body tilts. Since the hydraulic tilt table was only capable of rotating 20° in each direction, animals were initially held 20° head down, and then repositioned to 20° head-up while unit activity was recorded, so the change in body position was 40° (see Fig. 1). The 40° tilt occurred over a period of 16 sec (2.5°/sec); this slow tilt velocity was required to assure that units were not lost. During recordings, the room was darkened. An array of 10 LED lights, which generated a light intensity of 300 lumens, was placed 28 cm in front of the animal’s face. The LED light array was computer-controlled and illuminated for 2 seconds, from 8-10 sec prior to each 40° rotation (see Fig. 1). Since the area surrounding the tilt table was dark, the light cue was presumably highly salient. The light cue was provided prior to tilts throughout the acclimation period, such that animals experienced hundreds of cued tilts before the onset of neuronal recordings. During all trials, the animal was monitored to ensure that it was awake, and that it did not exhibit any indicators of distress.

**Figure 1.**
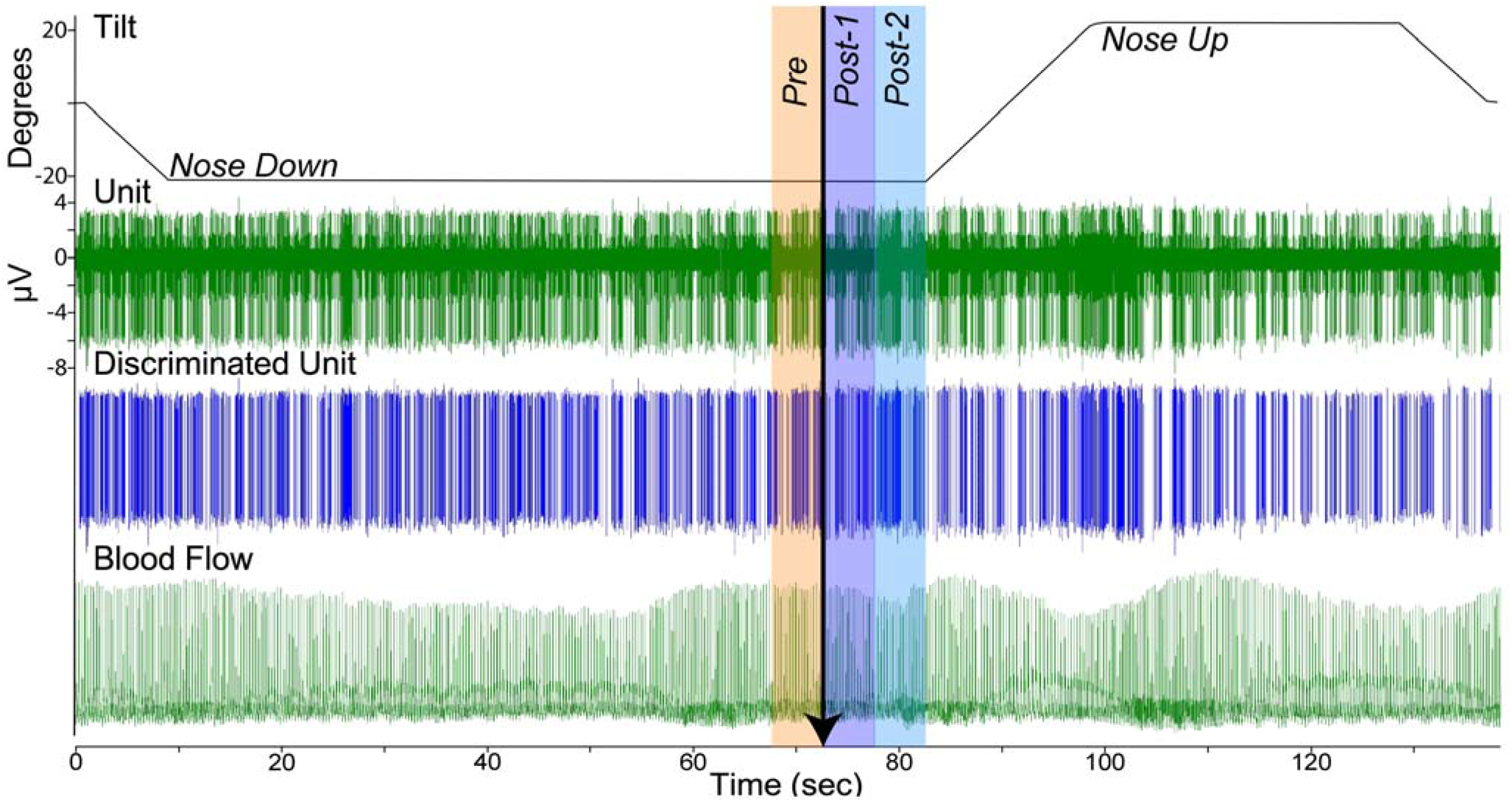
Procedures and data collected during a trial. Animals were positioned 20° nose down for a variable time period, a light cue was provided (onset indicated by arrow) that persisted for 2-sec, and a 40° head-up tilt (from 20° nose-down to 20° nose-up) was initiated at 10 sec after the onset of the light cue. Animals were maintained nose-up for 30 sec prior to being returned to the prone position. The top trace is a recording of table position, the second is a microelectrode recording of unit activity, the third shows discriminated activity for one unit, and the bottom trace is a recording of carotid artery blood flow. To determine if a unit’s activity was affected by the light cue, data collected during 2 time periods were compared using a paired t-test: 5-sec before light onset (Pre) and 5-10 sec after the light onset (Post-2).

After animals were well-adapted to head and body restraint and tilts, microelectrode penetrations into the brainstem were initiated. Epoxy-insulated tungsten microelectrodes (5MΩ, Frederick Haer, Bowdoin, ME) were maneuvered by using an x-y positioner and hydraulic microdrive (model 650, David Kopf Instruments, Tujunga, CA), which were attached to the recording chamber. Recordings were typically performed every weekday over a 4-5 month period (~100 electrode penetrations per animal). As in previous experiments, the ventral respiratory group (which is positioned dorsal to the RVLM) was used as a physiological landmark (4, 12). Activity recorded from brainstem neurons was amplified by a factor of 10,000 and filtered with a band pass of 300–10,000 Hz. The output of the amplifier was sampled at 25,000 Hz using a Micro1401 mk 2 data collection system and Spike2 version 7 software (Cambridge Electronic Design, Cambridge, UK). In addition, the ECG signal was amplified by a factor of 10,000 and filtered with a bandpass of 10-10,000 Hz and sampled at 2,500 Hz, whereas a voltage proportional to instantaneous carotid artery blood flow was generated by a Transonic Systems TS420 perivascular flow module and sampled at 100 Hz. Voltages produced by potentiometers mounted on the tilt table were sampled at 100 Hz to document table position. An example of data recordings is provided in Fig. 1.

When a putative RVLM unit was isolated (based on stereotaxic coordinates and positioning ventral to the cluster of neurons presumed to be the ventral respiratory group), the animal was rotated 20° head-down. The animal was held in this position for 1-4 minutes while baseline recordings were collected to determine if the unit exhibited CRA. Since the length of the head-down tilt period varied between trials, animals could not predict the occurrence of the light cue and subsequent head-up tilt. After the head-up tilt, the animal was maintained in the head-up position for 30 sec prior to being returned to the prone position (see Fig. 1). Trials were repeated as long as stable recordings from the unit could be maintained, with a maximum of ten trials per unit.

We also recorded the responses of many units to 10° (peak-to-peak) 0.5 Hz sinusoidal rotations in the pitch plane. These data were acquired to confirm that such small amplitude rotations do not elicit modulation of the activity of RVLM neurons, as reported previously (12).

### Data Analysis Procedures

The spike detection and sorting feature of the Spike-2 software was employed to segregate the activity of each unit in the recording field; only one or two units were typically present. Spike shape was considered across the duration of a trial as well as between trials, to ensure that data were consistently sampled from the same unit. Data were discarded if there was any doubt that activity was continually recorded from the same neuron. Subsequently, event markers were generated to indicate each action potential generated by a particular neuron; these event markers were used for subsequent analyses. In addition, event markers were placed at the peak of ECG R-waves and peak blood flow recorded from the carotid artery, to allow a determination of heart rate.

Each data file was reviewed to determine if animal movement was present, as indicated by the presence of electromyographic activity on ECG recordings (see Fig. 2), and if the activity of the unit changed systematically before or during the periods of movement. If movement persisted for only a few seconds, as in Fig. 2, the segment containing movement was cropped from the record using Spike-2 software. If movement was prevalent during the trial, it was discarded, but units were typically lost when substantial movement occurred.

**Figure 2.**
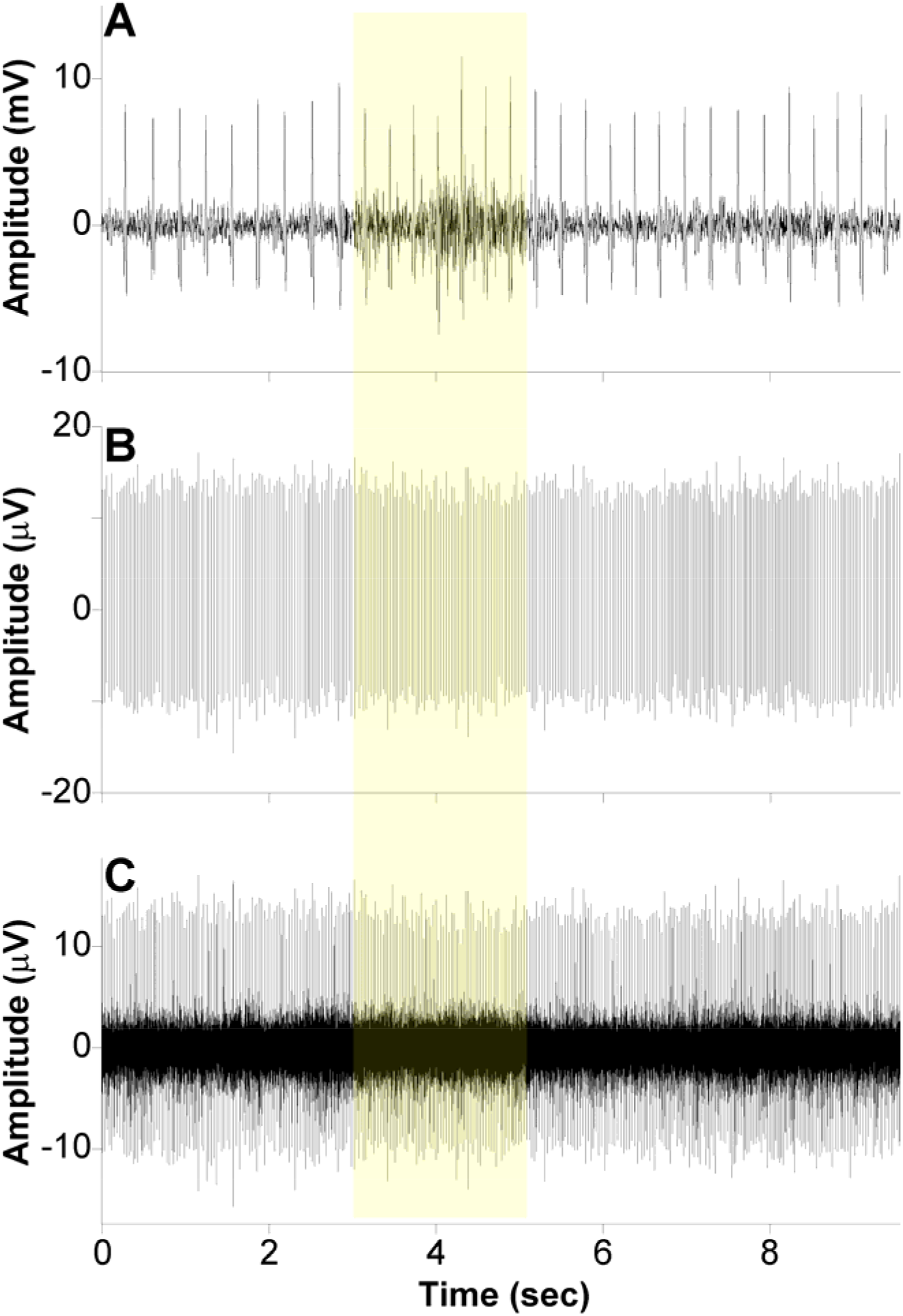
Recording of unit activity during a brief period of animal movement reflected by electromyographic activity on the ECG recording. **A**: ECG recording; **B:** discriminated activity of one unit; **C:** microelectrode recording of unit activity. Shading highlights the period during which electromyographic activity was detected.

#### Detection of CRA

Procedures similar to those in our prior studies were used to determine if a unit exhibited CRA (4, 12). Continuous data segments recorded while the animal was tilted head-down, without electromyographic activity evident on the ECG recording (indicating animal movement; see Fig. 2A), were used for analyses. Trigger pulses coincident with the R-wave of the ECG or the peak of the carotid blood flow signal were used for the construction of post-R wave (or post- carotid flow) interval histograms (10-ms bin width) of unit activity (Datapac software, Run Technologies; Mission Viejo CA). A histogram of intervals between consecutive R waves was simultaneously constructed. The trigger pulses were also used to construct an average of the carotid blood flow signal. These histograms and averages were constructed using data segments of varying length (30 – 160 s). As noted above, multiple trials were often conducted for each unit, and a separate analysis was performed for each trial. A ratio of peak-to-background counts was calculated for each histogram. To classify a neuron as having CRA, the ratio of peak-to-background counts in the histogram had to exceed a value of 1.5 in the majority of runs for each unit. Fig. 3 shows histograms illustrating CRA in two trials for one unit.

**Figure 3.**
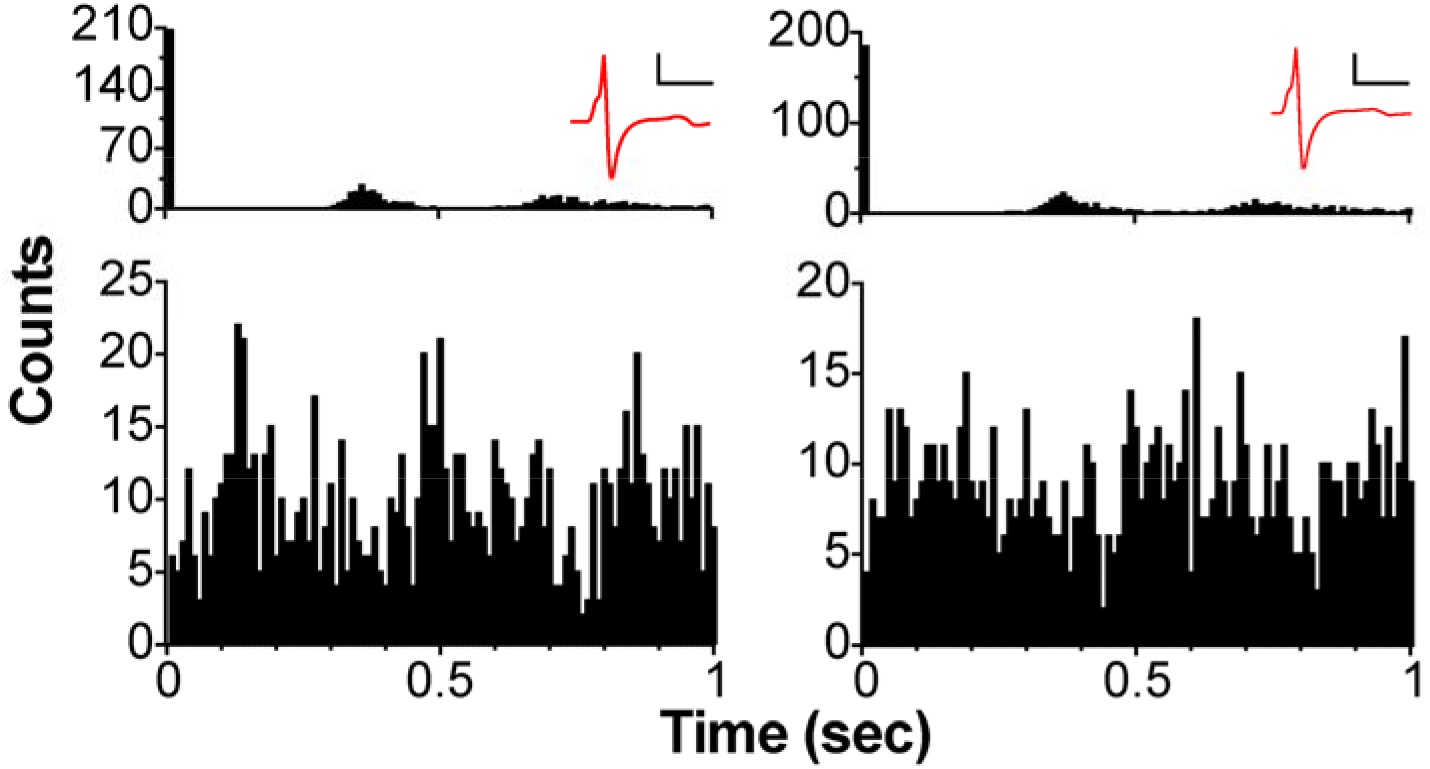
CRA evident in two trials for an RVLM unit. Each column shows histograms triggered by ECG R-waves of intervals between consecutive R-waves *(top)* and intervals between action potentials recorded from the unit *(bottom).* Bin width is 10 msec. The time period shown encompasses three cardiac cycles. In the top panels, there was a sharp peak at time 0 (trigger), and the subsequent peaks are less sharp due to variability in the cardiac cycle. In the bottom panels, it is evident that unit activity is related to the cardiac cycle. The red waveform at the top of each column is the averaged action potential for the trial. In both trials, the spike shape is similar showing that the same unit was recorded throughout. Scale bar for action potential: x-axis depicts 1 msec; y-axis shows 5 mV.

#### Reponses to Light Cue and 40° Tilts

Subsequent analyses were performed using Matlab software (Mathworks, Natick, MA). Unit firing rate and heart rate were determined when the animal was in the head-up position, as well as during a similar time period prior to the light cue when the animal was positioned head-down. In addition, unit firing rate and heart rate were ascertained for the 5-sec period prior to the light cue and during the two subsequent 5-sec periods prior to tilt onset (see Fig. 1). Since the light cue persisted for 2 sec, the light was illuminated for the first two seconds of the Post-1 period indicated in Fig. 1. Data for units with five or more 40° tilt trials were considered in subsequent statistical analyses conducted using Prism 8 software (Graphpad software, San Diego, CA). Confidence intervals are reported as means ± one standard deviation.

#### Reponses to Sinusoidal Rotations

Our procedures for analyzing responses to sinusoidal rotations have been described in many publications (12, 37, 39, 40, 42, 43). Neural activity recorded during sinusoidal rotations was binned (500 bins/cycle) for ~35 stimulus repetitions and fitted with a sine wave with the use of a least-squares minimization technique. Averaged data were inspected during recording sessions and trials were repeated when a unit appeared to respond to the rotation. Two major criteria were used to determine if a unit’s activity was modulated significantly by rotations: a signal-to-noise ratio > 0.5 and only one prominent first harmonic. We also considered whether responses were consistent between trials, to ensure that apparent responses were not a reflection of a unit’s rhythmic spontaneous firing rate near the frequency of the rotation. If rhythmic activity was present, the timing of the peak of the apparent response in relation to stimulus would change appreciably between trials. In the few cases where this type of rhythmic activity was present, the data were discarded.

#### Reconstruction of unit locations

After data recording was concluded in an animal, electrolytic lesions were made at defined coordinates by passing a 100-μA negative current for 60 s through a 0.5-MΩ tungsten electrode. Subsequently, animals were deeply anesthetized using an intramuscular injection of 20 mg/kg ketamine and 0.2 mg/kg acepromazine, followed by an intraperitoneal injection of 40 mg/kg pentobarbital sodium, and then perfused transcardially with saline followed by 10% formalin. The brain was removed and postfixed in 10% formalin for at least two weeks prior to being sectioned transversely at 50-μm thickness using a freezing microtome. Sections were mounted serially on slides and stained using thionine. Images of each section were acquired using a Nikon Eclipse E600 microscope equipped with a Spot digital camera and software (Spot Imaging, Sterling Heights, MI). Photomontages of sections were generated using Adobe Photoshop software (Adobe Inc., Mountain View, CA) and used to map unit locations. Unit locations were reconstructed with respect to the position of lesions, the coordinates of each track, and depths of recordings.

## RESULTS

Activity was recorded from 250 units with CRA whose location was histologically confirmed in the RVLM: 142 neurons in one animal and 108 in the other. Unit activity was collected from the animals over a period of 134-147 days. In addition, responses to tilts were recorded for an additional 92 units (56 in one cat and 36 in the other) without CRA that were interspersed with the neurons whose firing pattern was correlated with the cardiac cycle.

Statistical analyses compared responses across 40° tilt trials for each unit (e.g., compared unit firing rate when the animals were in nose-down and nose-up positions). Recordings were stable (e.g., spike shape remained consistent) for 55 RVLM units with CRA during 5 or more trials (5 trials for 19 units, 6 trials for 12 units, 7 trials for 8 units, 8 trials for 5 units, 9 trials for 8 units, and 10 trials for 3 units). The locations of these units are indicated in Fig. 4. Additionally, statistical analyses were conducted for 36 RVLM units without CRA whose responses during 5 or more trials were collected.

**Figure 4.**
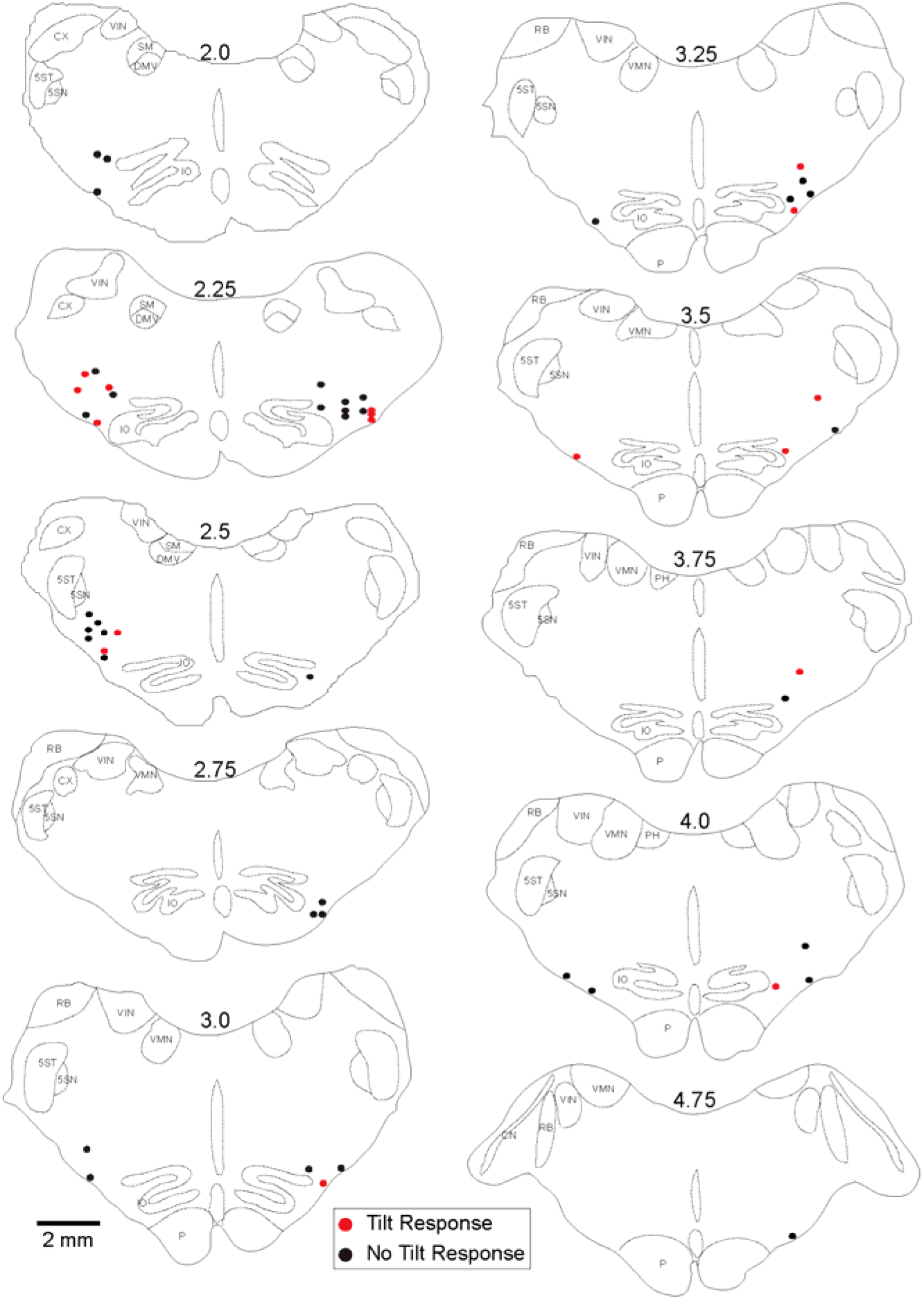
Locations of RVLM neurons with CRA whose responses to tilts were recorded during 5 or more trials. The numbers above each section indicate distance in mm rostral to the obex. Abbreviations: 5SN, spinal trigeminal nucleus; 5ST, spinal trigeminal tract; CN, cochlear nuclei; CX, external cuneate nucleus; DMV, dorsal motor nucleus of the vagus; IO, inferior olivary nucleus; P, pyramid; PH, prepositis hypoglossi; RB, restiform body; SM, medial nucleus of the solitary tract; VIN, inferior vestibular nucleus; VMN, medial vestibular nucleus.

### Rhythmic activity of units

Fig. 5 compares the peak and trough activity for the 55 units with CRA tested with multiple trials. Fig 5A shows the counts (number of action potentials) in the peak and trough periods, while Fig. 5B indicates the ratio of the peak and trough counts. Each point represents the average counts across all trials for a particular unit. For every unit tested using 5 or more trials, a paired t-test revealed that the average peak counts were significantly greater (P<0.05) than the average trough counts. P values ranged from <0.0001 to 0.02, with most units having P<0.003, and only 4 having P values between 0.01 and 0.05. On average, peak counts were 2.5±1.0 (SD) times greater than trough counts. Although CRA was present across trials for units that showed this activity, the ratio of peak to trough counts could vary between trials, as shown in Fig. 6 for 8 units, as well as the examples in Fig. 3.

**Figure 5.**
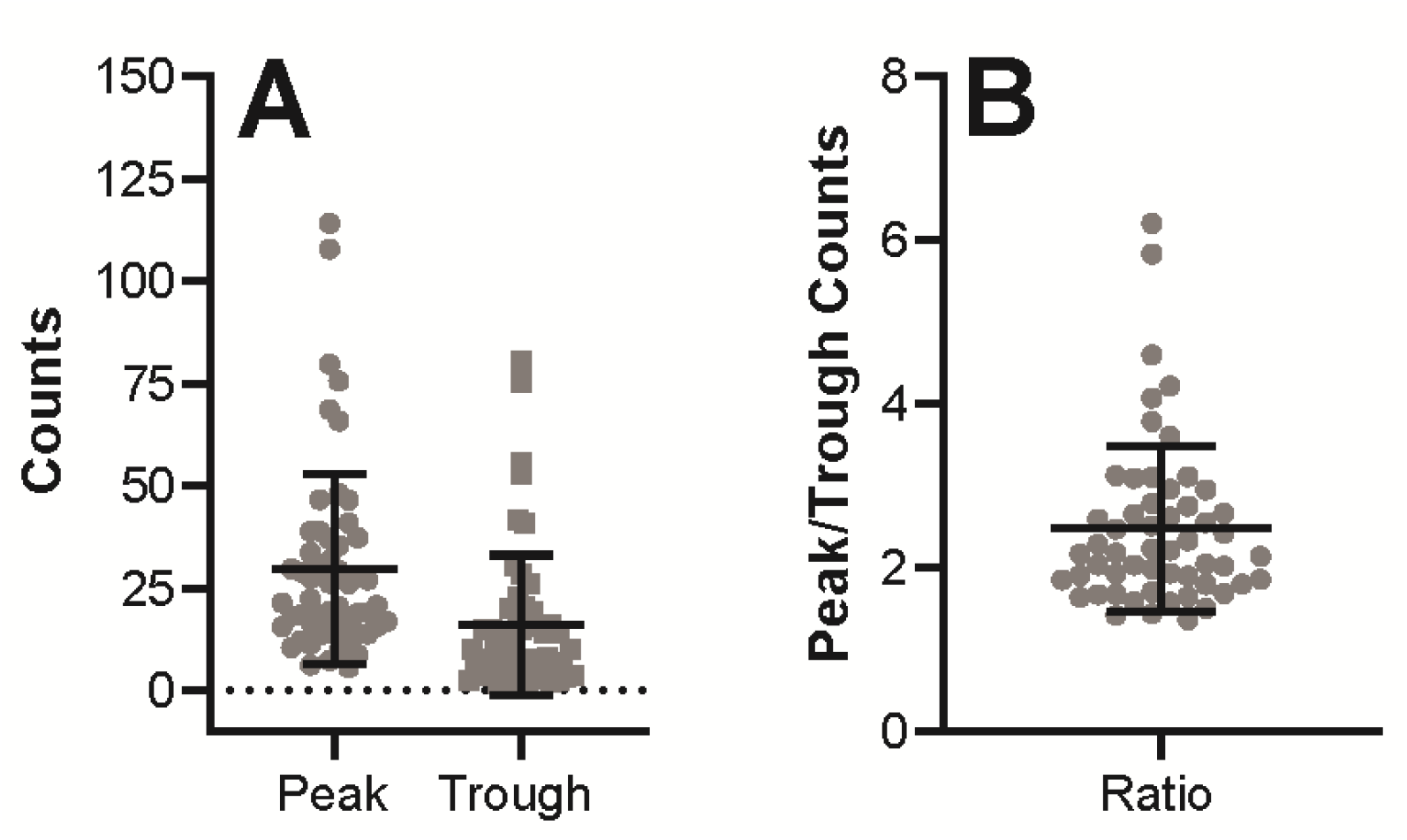
A: Plot of counts during peak and trough periods of cardiac-related activity for each of the 55 units with CRA. B: Ratio of peak and trough counts for each of the 55 units. Error bars designate mean ± one standard deviation.

**Figure 6.**
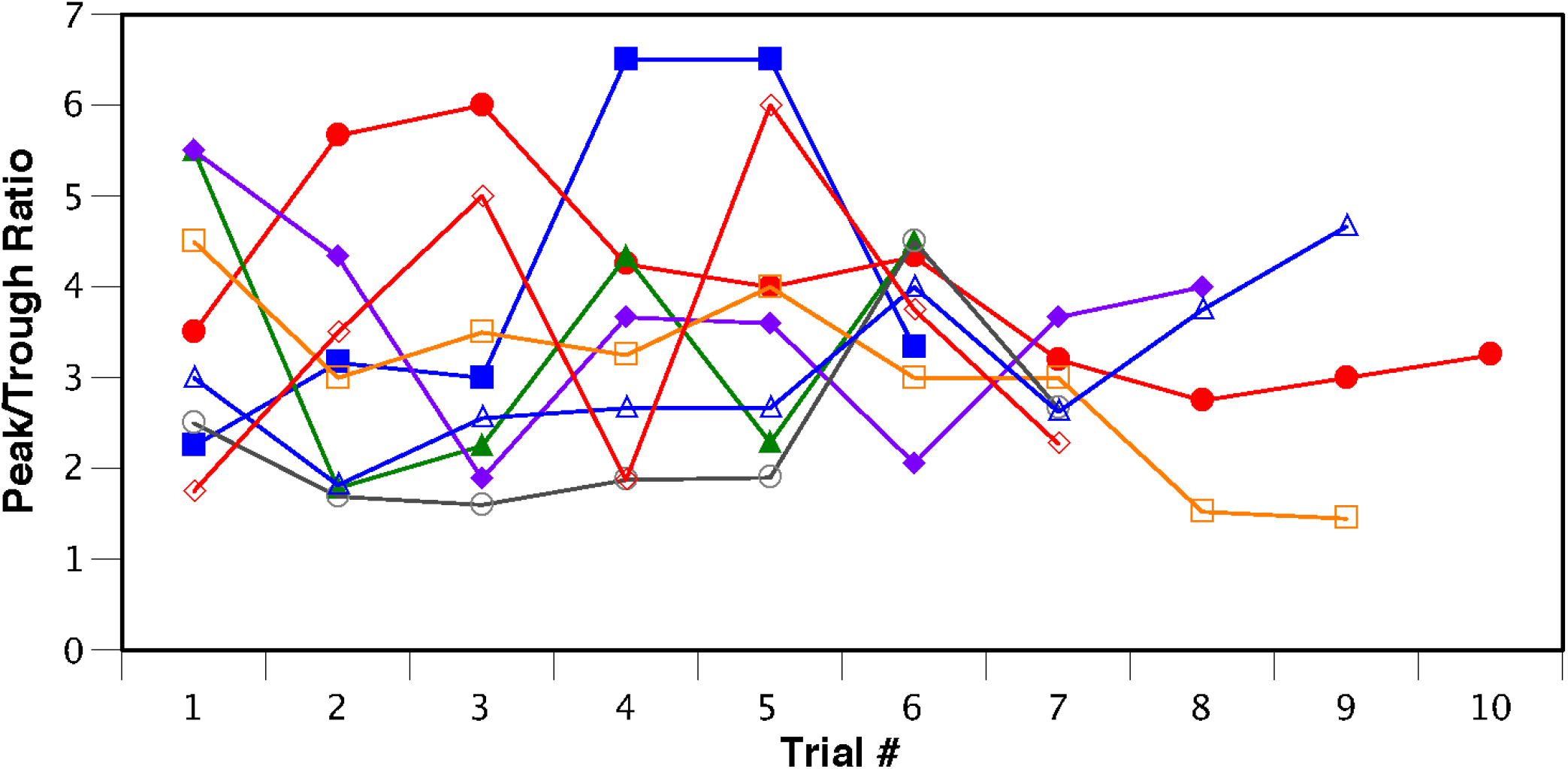
Ratio of peak to trough activity related to the cardiac cycle for 8 units; data for each unit is designated by different symbols and colors. Although the units exhibited CRA during each trial, the magnitude of the responses (as reflected in the peak/trough ratio) varied during the course of recording.

**Figure 7.**
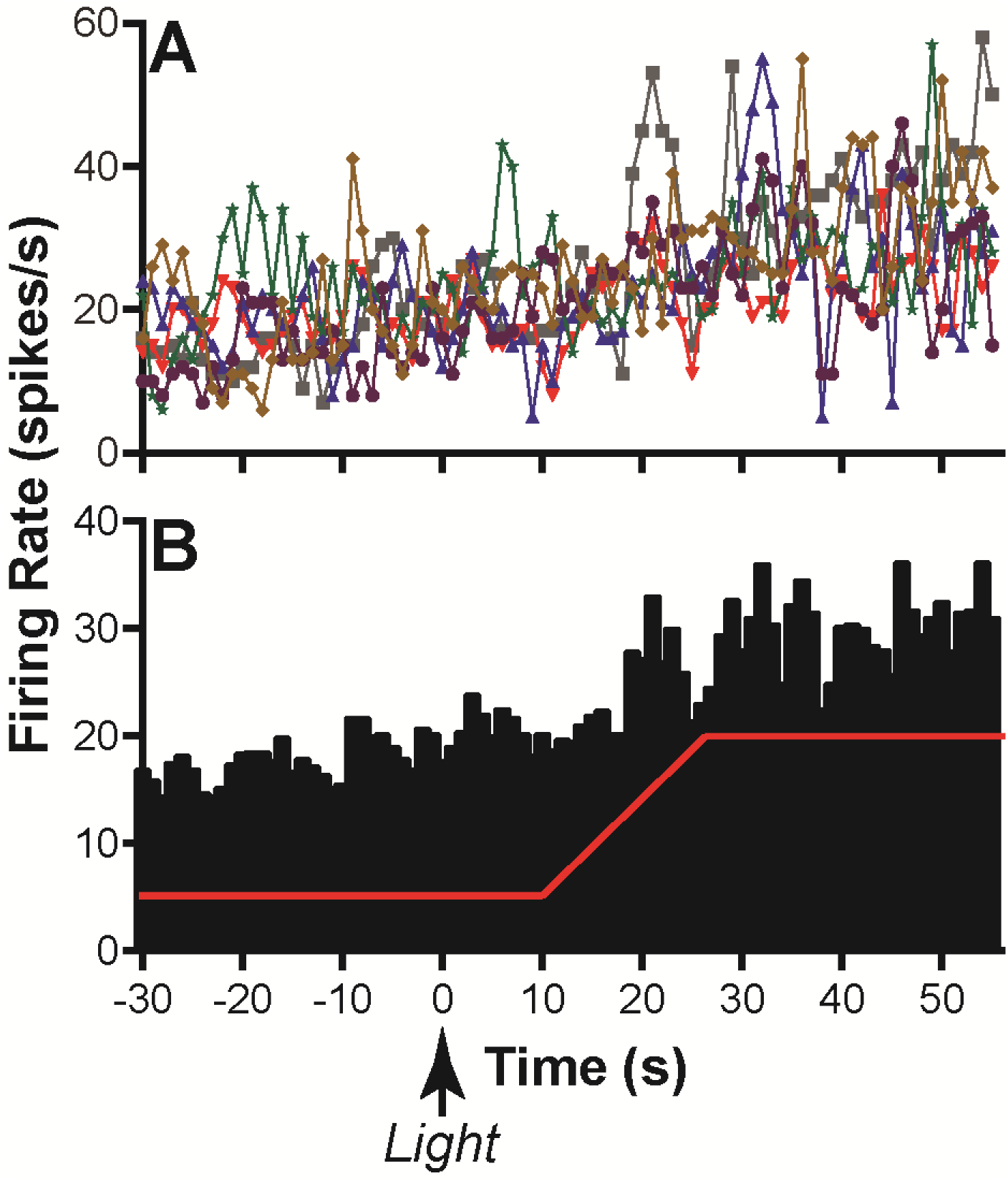
Change in activity for one RVLM unit with CRA during 40° head-up tilt. **A**: Unit activity during 1-sec bins for 6 individual trials; data for each trial is depicted by different symbols and colors. **B:** Average activity during each 1-sec bin for the 6 trials. A red line indicates the 40° head-up tilt (transition from 20° nose-down to 20° nose-up). An arrow designates the onset of the 2-sec light cue, which preceded each tilt.

**Figure 8.**
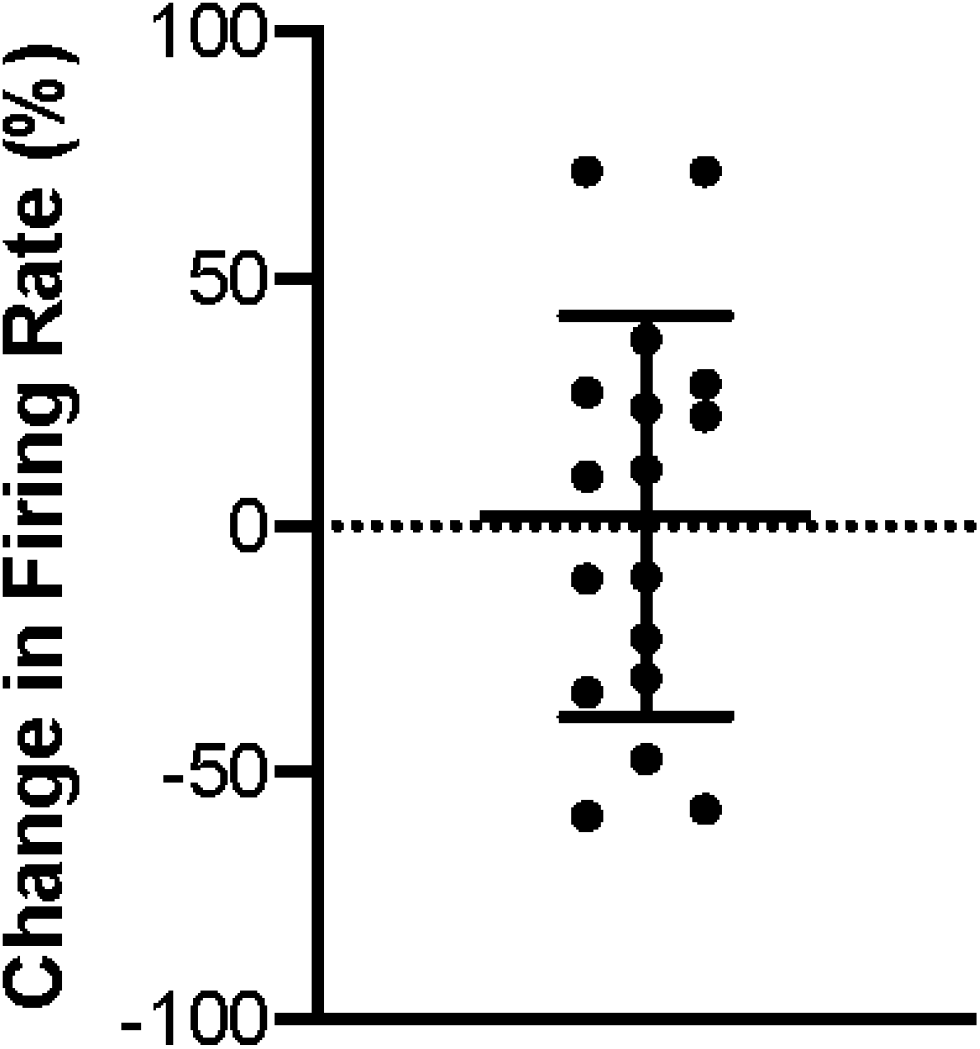
Percent change in firing rate during 40° head-up tilts (transition from 20° nose-down to 20° nose-up) for 17 RVLM units with CRA that had significant responses to changes in body position. Error bars indicate mean ± one standard deviation.

### Responses to Tilt

Activity of RVLM units with CRA was compared when animals were in the 20° nose-down and 20° nose-up positions. Fig. 7 shows the change in activity for one unit during the 40° head-up tilt (change in position from 20° nose-down to 20° nose-up). Fig. 7A indicates the unit’s activity during 1-sec bins for 6 individual trials, whereas Fig. 7B shows the average activity in each bin across trials. For the 55 units with 5 or more trials, we compared neuronal activity when the animals were nose-down and nose-up using a paired t-test. For 17 of these units (31%), activity was significantly different (P<0.05) when the animals were nose-down and nose-up. Fig. 8 indicates the percentage difference in firing rate during the nose-up position relative to the nose-down position of these cells. When the analysis was limited to units with 6 or more tilt repetitions (and thus greater statistical power), a paired t-test revealed that 14/36 cells (39%) responded significantly to head-up tilts.

The RVLM units with CRA whose firing rate was significantly altered by 40° changes in body position were intermingled with those whose firing rate was unaffected by the tilts, as shown in Fig. 4. Neurons that responded to tilts were located at 3.7±0.5 mm rostral to the obex and 3.4±0.7 mm lateral to the midline, while unresponsive units were located at 3.6±0.6 mm rostral to the obex and 3.3±0.5 mm lateral to the midline. A two-way ANOVA showed that the distances from the obex (P=0.97) and from the midline (P=0.93) were not significantly different for RVLM neurons whose activity was modulated or unaffected by 40° head-up tilts.

Fig. 9 shows the average percent change in activity in the nose-up position for the 17 RVLM units with significant responses to tilt (e.g., nose-up activity/nose down activity * 100%). The activity of 9 units increased in the nose-up position (relative to the nose-down position), while the activity of 8 units decreased. The average absolute firing rate of these units when animals were positioned 20° head-up differed 34±21% from that when animals were tilted 20° head-down.

**Figure 9.**
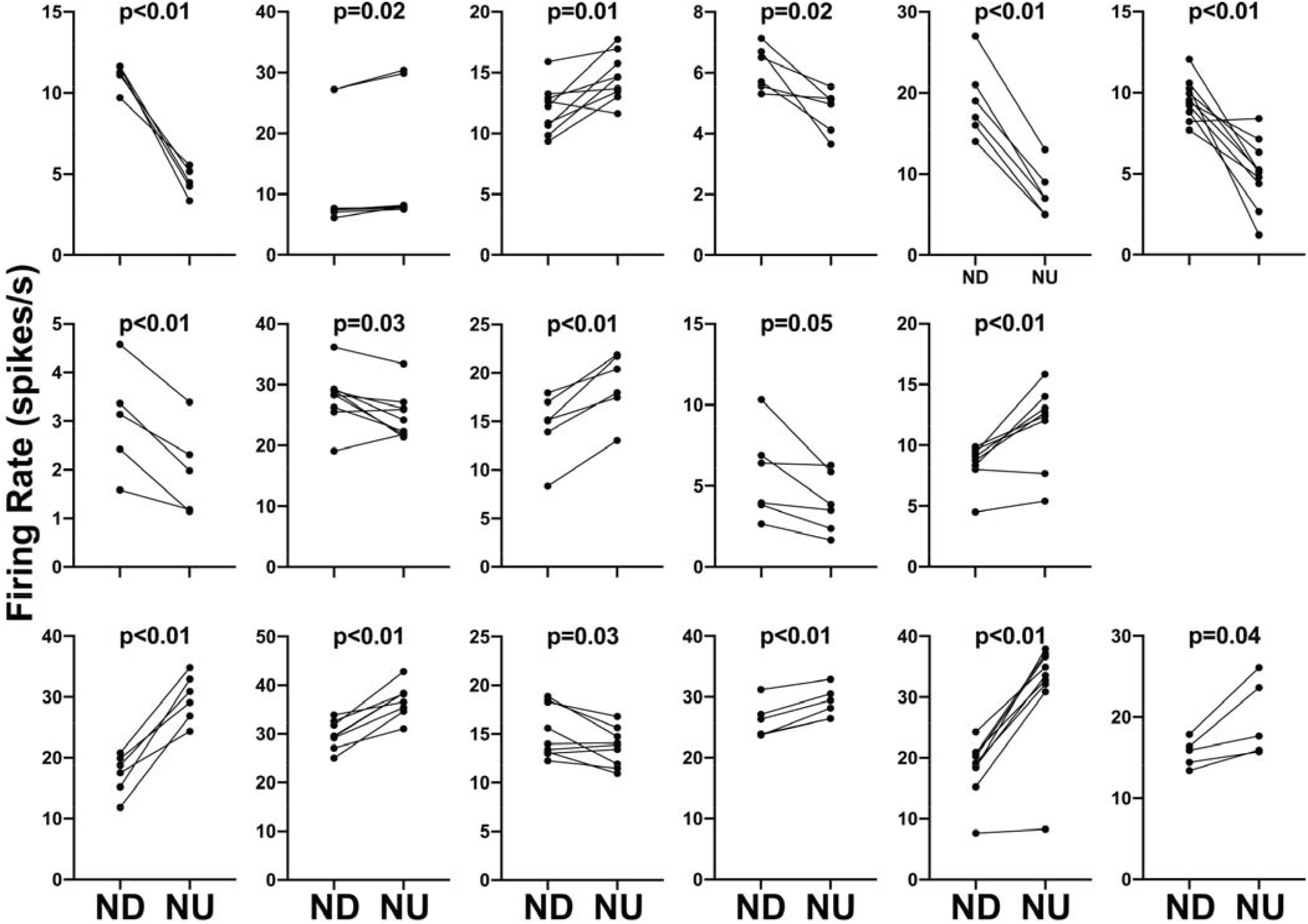
Firing rate during each trial for the 17 RVLM units with CRA that responded significantly to head-up tilt. Each panel compares the firing rate for an individual neuron when the animal was positioned 20° nose-down (ND) and 20° nose-up (NU). P values above each panel indicate the significance of the difference in firing rate in the ND and NU positions (two-tailed paired t-test). The greatest significance level indicated is p<0.01 (i.e., responses that differed at p<0.001 are indicated as p<0.01).

Responses were additionally recorded for the 17 units during 10° sinusoidal rotations in the pitch plane at 0.5 Hz. Only 2 of these units had significant responses to these rotations in accordance with established criteria (12, 37, 39, 40, 42, 43). Responses to 10° sinsusoidal rotations for the two units are illustrated in Fig. 10. Only 1 of the 8 units with large (>30%) changes in activity during 40° head-up tilts responded significantly to 10° sinusoidal rotations. None of the units that did not respond to 40° trapezoidal rotations had activity that was modulated by 10° sinusoidal tilts.

**Figure 10.**
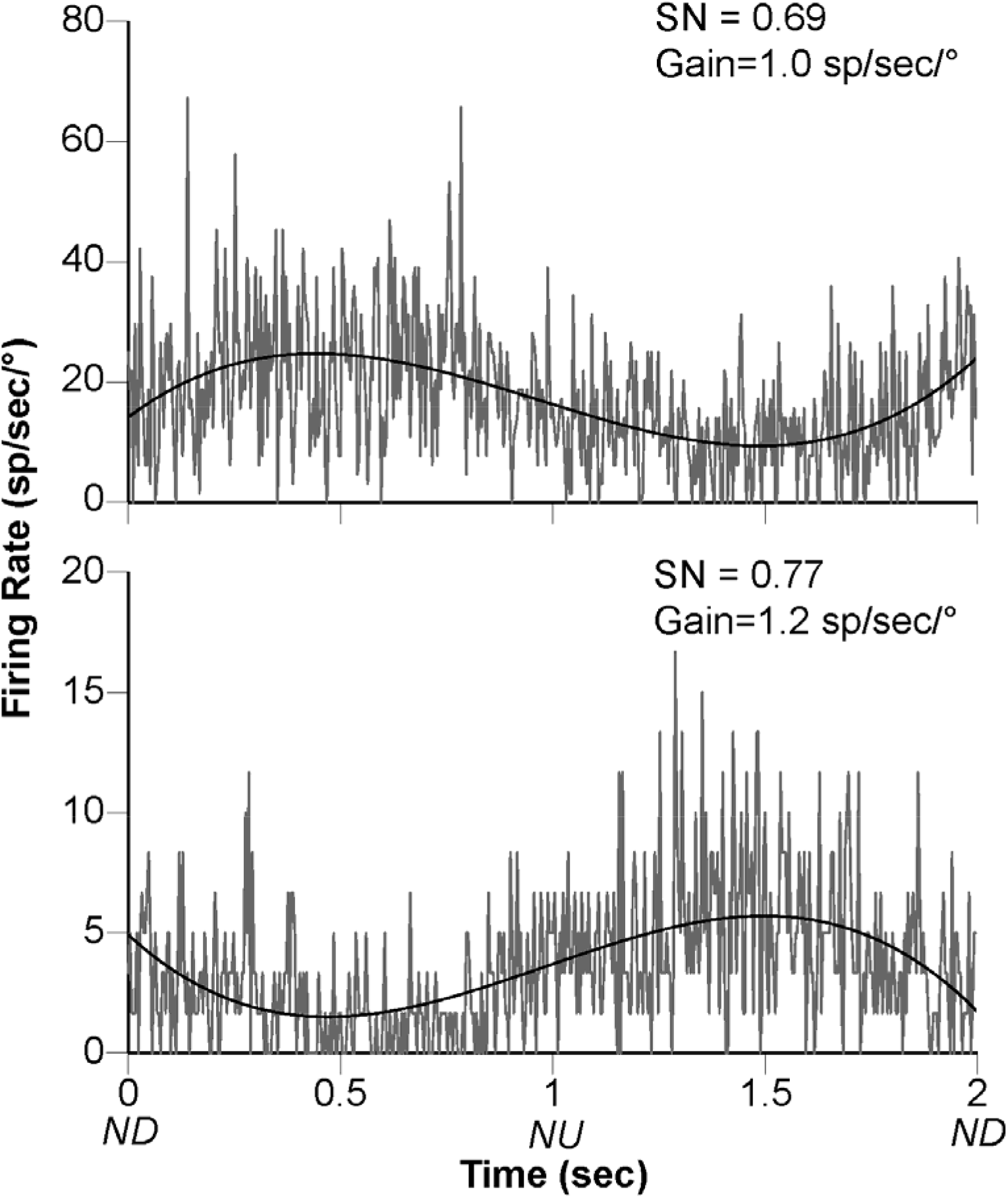
Responses to 10° sinusoidal rotations in the pitch plane at 0.5 Hz for the two RVLM neurons with CRA that responded to the stimulus. Gray traces show averaged activity during ~30 stimulus repetitions; the bin width was 4 msec (500 bins/trace). Black lines show the sine wave fit to each response. The signal-to-noise (SN) ratio and gain for each response are indicated. Abbreviations: ND, nose-down; NU, nose-up.

We also considered whether 40° changes in body position altered the firing rate of RVLM neurons without CRA. Of 36 cells tested with 5 or more tilt repetitions, 9 (25%) responded significantly to head-up rotations. P values ranged from 0.0006 to 0.04, with 4 responses having P<0.01. Activity increased in 6 units during head-up tilts and decreased in the other 3. The average absolute change in firing rate for the units during 40° head-up tilts was 34±42%. A two-sided Fisher’s exact test failed to show that the fraction of units that responded to 40° rotations differed for RVLM units with and without CRA (P=0.64).

To ascertain whether baroreceptor inputs might be responsible for responses of RVLM neurons to 40° rotations, we additionally monitored heart rate changes during head-up tilts. Only 6/17 units that responded to 40° tilts and 6/38 that failed to respond to 40° tilts exhibited significant (P<0.05) changes in heart rate during the rotations. In 10/12 of these cases, heart rate increased during head-up tilts, but for two units head rate decreased during the rotations. On average, heart rate increased by 2.4±3.8% during head-up tilts that elicited a significant change in the firing rate of RVLM neurons, and heart rate increased 1.1±2.2% during trials where units failed to respond to the rotations. The changes in heart rate during rotations that did or failed to alter the firing rate of RVLM neurons were not significantly different (P=0.13, two-tailed t-test). Fig. 11 compares the change in heart rate and unit firing rate that occurred during 40° tilts. A linear regression analysis failed to demonstrate a significant relationship between the change in heart rate and the change in unit activity resulting from head-up tilts (P=0.83; R^2^<0.001). When the analysis was limited to units with significant responses to tilts, the relationship between the change in heart rate and unit activity was also not significant (P=0.24; R^2^=0.09). Moreover, for the 8 units with large (>30%) changes in activity during head-up tilts, only one exhibited a significant change in heart rate during the rotations.

**Figure 11.**
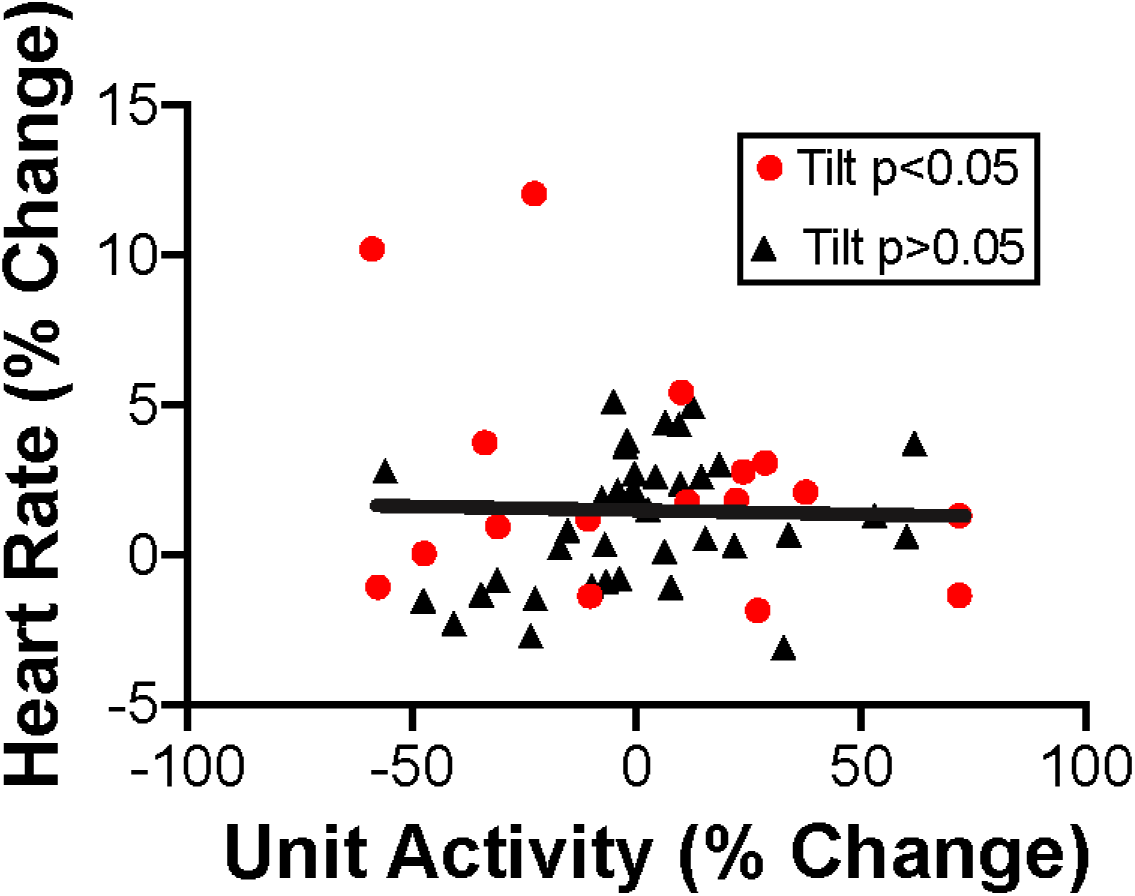
Comparison between the change in firing rate of RVLM units with CRA and heart rate that occurred during 40° head-up tilts. Red symbols designate the units with significant responses to head-up tilts (P<0.05, comparison of activity in the 20° nose-down and 20° nose-up positions using a paired t-test). A solid line shows a best-fit of the data from a linear regression analysis (P=0.83; R^2^<0.001).

All runs were examined carefully for changes in electromyographic activity on the ECG trace (see Fig. 2) that could have reflected stiffening of the body or movement during whole-body tilts. Although transient occurrences of EMG activity were noted, unit activity did not appreciably change during these periods, as exemplified in Fig. 2.

### Preparatory responses prior to tilts

A light cue preceded each tilt by 10 sec, and we evaluated whether activity of RVLM units changed after the light cue and prior to tilts. Fig. 12 shows that only four of the 55 RVLM units with CRA had significant changes in activity (p<0.05, two-tailed paired t-test) in the 5-second period prior to tilts (Post-2 period indicated in Fig. 1) compared to the 5-sec period prior to the light cue (Pre- period indicated in Fig. 1). P values were 0.01 for one cell and between 0.04 and 0.05 for the others. Only 1 of the 17 units with significant responses to head-up tilts exhibited a significant change in firing rate prior to the rotation. Of the 36 cells without CRA tested in multiple trials, only 1 had significant changes in firing rate in the pre-light and post-light 2 periods indicated in Fig. 1. The firing rate of this unit was not significantly altered by head-up tilt.

**Figure 12.**
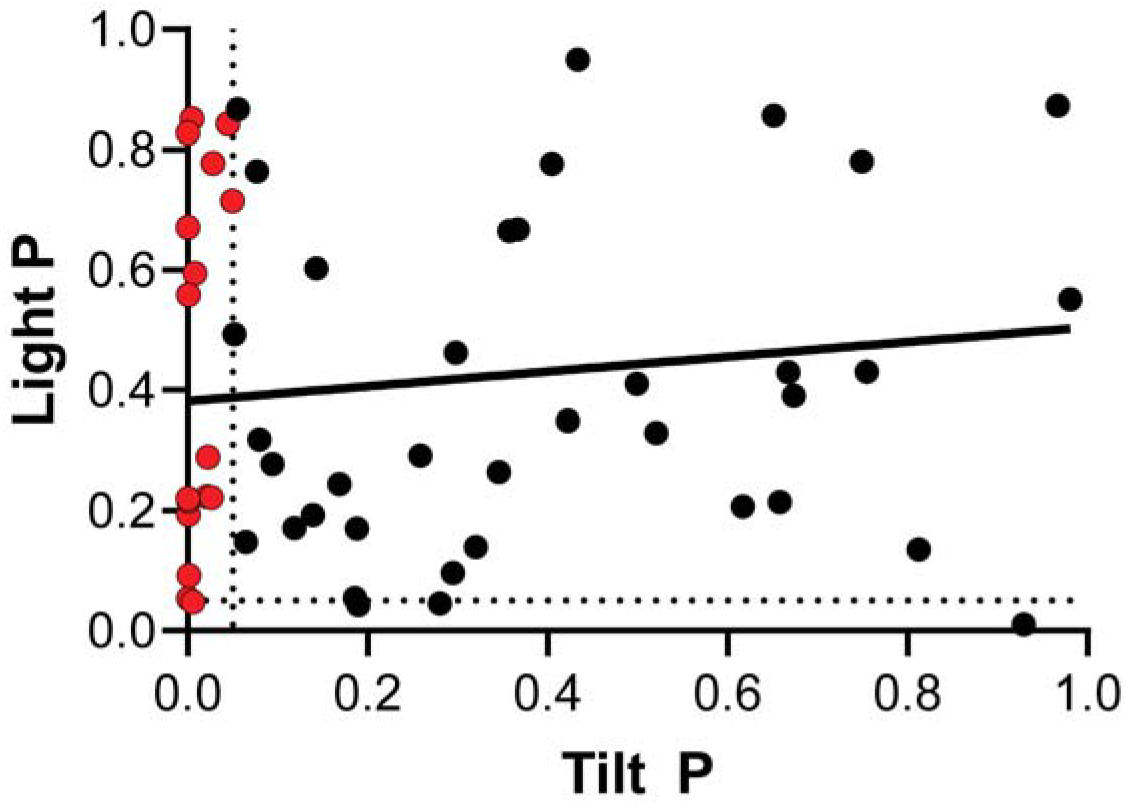
P values (two-tailed paired t-test) for responses to 40° head-up tilts (comparison of activity when animals were positioned 20° nose-down to that when animals were positioned 20° nose-up) as well as responses to the light cue prior to tilts (comparison of activity in the Pre- and Post-2 periods illustrated in Fig. 1). Data for the 17 RVLM units with CRA with significant responses to head-up tilts (P<0.05) are indicated by red symbols. A solid line shows a best-fit of the data from a linear regression analysis. This analysis showed there is not a strong relationship between the significance of responses to tilt and the significance of responses to the light cue (P=0.34; R^2^=0.02). Dashed lines indicate P=0.05 for responses to tilts and the light cue.

## DISCUSSION

This main finding of this study is that large (40°) head-up rotations result in a significant change in activity of an appreciable fraction of RVLM neurons of conscious felines. It was also verified that 10° sinusoidal rotations that robustly modulate the activity of neurons in other brainstem areas that process vestibular signals (37, 39, 42) have little effect on RVLM unit activity in conscious animals, corroborating findings in a previous study (12). In contrast, neurons in brainstem regions that provide inputs to the RVLM, including the caudal vestibular nuclei (2, 42) and lateral tegmental field (39, 44), exhibit robust responses to small-amplitude sinusoidal rotations in both decerebrate and conscious animals. The amplitude of movement required to alter the activity of an appreciable fraction of RVLM neurons was much larger in conscious cats. The neural mechanisms responsible for establishing a threshold of movement amplitude required to alter the firing rate of RVLM neurons in conscious animals remains to be determined. The midline cerebellum has a well-established role in modulating the responsiveness of brainstem neurons to vestibular stimuli, and the cerebellar uvula (lobule IX) provides multisynaptic inputs to RVLM neurons (47–49). It remains to be determined whether the cerebellar uvula, or other brain areas, adjust the responsiveness of RVLM neurons to vestibular stimuli. However, it is evident that supratentorial brain areas are critical for establishing a threshold amplitude of body movement required to alter the activity of RVLM neurons, since they respond to small-amplitude tilts in decerebrate but not conscious animals (12).

The neurons sampled in this study were located ventral to a cluster of neurons with respiratory activity in the ventrolateral medulla (presumably the ventral respiratory group). Most neurons in this area had CRA, suggesting that they participated in regulating SNA (5), and their locations were confirmed in the brainstem area of felines that contains reticulospinal neurons that project to sympathetic preganglionic neurons (3, 11, 18, 36, 57). In the conscious preparation used in this study, it was not feasible to use antidromic stimulation to verify that the neurons sampled projected to the thoracic spinal cord. Nonetheless, based on the evidence discussed above, it is highly likely that pre-sympathetic neurons were targeted in these experiments.

RVLM neurons are inhibited by baroreceptor inputs (3), and thus one possibility is that responses to the large-amplitude pitch rotations used in this study were secondary to unloading of baroreceptors during head-up rotations. However, there was little correlation between changes in SNA, as gauged by alterations in heart rate, and changes in unit activity during tilts. The robust responses to tilts of a group of neurons in this study were thus most likely due to activation of vestibular receptors.

Of the neurons whose firing rate was significantly altered by 40° head-up tilts, the activity of approximately half increased and the activity of the other half decreased. Several studies have demonstrated that unlike baroreceptor reflexes, VSR are anatomically patterned in felines, and differ in the upper and lower body (29–32, 61). Although the axons of some RVLM neurons branch to multiple thoracic and lumbar cord segments, others have projections to either the rostral or caudal portions of the intermediolateral cell column (3, 14, 18, 57). It is speculated that the RVLM neurons whose activity increased during 40° head-up tilts affected sympathetic outflow to the lower body (to produce vasoconstriction and minimize blood pooling), whereas those whose activity decreased affected sympathetic outflow to the upper body (to reduce vasoconstriction such that stable blood flow was maintained despite gravitational effects). The diversity of responses of RVLM neurons to head-up rotations is also further evidence that they were elicited by vestibular inputs and were not caused by baroreceptor unloading.

Intense muscle contraction (the muscle pressor reflex) can also affect the activity of RVLM neurons, at least in decerebrate and anesthetized animals (15, 28). Although the animals in this study were acclimated to remain sedentary during the experimental sessions, vestibulospinal reflexes likely resulted in alterations of the activity of the trunk muscles during head-up tilts (19). These changes in muscle activity were detected as electromyographic activity on ECG recordings but were not correlated with variations in RVLM neuronal firing rates. Thus, the muscle pressor reflex is unlikely to have triggered the alterations RVLM unit activity during head-up rotations.

Although units classified as having CRA exhibited this activity across runs, for some neurons the magnitude of cardiac-related responses varied between trials. A prior study provided some evidence that responses of RVLM neurons to baroreceptor inputs vary over time in conscious animals (4), and the present data validate these findings. It seems likely that factors such as vigilance and stress could alter the excitability of RVLM neurons, as they receive inputs directly and indirectly from a variety of supratentorial brain structures, including insular and prefrontal cortex, the amygdala, and several hypothalamic nuclei (for review, see (5)). In addition, it is feasible that the responses of RVLM neurons to other signals, including vestibular inputs, could be altered during behavioral paradigms with high likelihood of unexpected postural alterations that affect blood distribution in the body. This prospect remains to be tested experimentally.

A light cue preceded each 40° head-up tilt, and we evaluated whether RVLM unit activity was altered following the cue and prior to the onset of rotations. There was no evidence that changes in activity of most units were initiated prior to the onset of rotations, including neurons with robust responses to tilts. These findings extend prior observations that changes in heart rate and cerebral blood flow do not precede 60° head-up tilts signaled by a light cue (46). Conversely, it is well-established that feedforward (preparatory) cardiovascular responses are associated with exercise (13, 59). For example, increases in muscle blood flow, blood pressure, and heart rate occurred in trained rats just after they were placed on a treadmill and prior to the onset of exercise (1). The simplest interpretation of these findings is that feedforward cardiovascular responses are associated with active movement, but not passive movements that require cardiovascular adjustments. However, there are a number of caveats to this conclusion. First, the magnitude of changes in SNA are larger during exercise (9) than during 40° postural alterations (61), such that feedforward cardiovascular adjustments may only occur in circumstances where large changes in SNA are needed. It is also noteworthy that these studies did not include a positive control to conclusively demonstrate that the animals associated the light cue with head-up rotations, although prior studies showed that cues such as those used in these experiments are highly salient without producing startle (8, 25, 35, 53). It seems highly likely that after hundreds of trials, the felines could associate the light cue in a darkened room with the ensuing tilt.

Several other limitations to these studies should be considered. The experiments had a similar design to classical neurophysiological studies in nonhuman primates, in that only two animals that were well-acclimated for experimental procedures were used, but responses of a large number of neurons were sampled in each animal over a period of several months (22, 27, 45, 50, 54, 58). Since the findings were consistent in both animals, it does not appear the data were appreciably biased by physiological differences between individuals. Hence, it is unlikely that different conclusions would be reached by repeating the studies in additional animals. As noted above, although there are many advantages to using conscious animals in neurophysiological studies, a limitation is that their vigilance can vary over time, altering responses to stimuli. We addressed this limitation by acclimating the animals to the experimental paradigm for several weeks prior to the onset of data collection, and by carefully observing animals during recording sessions to assure they were awake.

### Perspectives and Significance

This is one of the first studies to examine in conscious animals the activity of brainstem neurons that presumably participate in the regulation of SNA and blood pressure. The findings indicate that an appreciable fraction of RVLM neurons with CRA respond to large head-up postural alterations but not to small changes in body position that robustly activate vestibular receptors and neurons in other brainstem regions, including those that provide inputs to the RVLM (37, 39, 42). The vestibular system evolved to detect very small changes in head position (17), but VSR are only required during large changes in body position in space that affect blood distribution (61, 65). Since RVLM neurons respond to small-amplitude rotations in decerebrate animals (12, 63), our data suggest the existence of supratentorial brain circuitry that suppresses vestibular influences on the activity of RVLM neurons and SNA unless these inputs are physiologically warranted. Further research is needed to decipher the neural mechanisms through which the response suppression occurs, and if the suppression varies depending on behavioral context.

## ACKNOWLEDGEMENTS

The authors thank Krista Roszkowski for contributing to data collection in these experiments.

